# Improved N- and O-Glycopeptide Identification using High-Field Asymmetric Waveform Ion Mobility Spectrometry (FAIMS)

**DOI:** 10.1101/2022.12.12.520086

**Authors:** Kathirvel Alagesan, Rina Ahmed-Begrich, Emmanuelle Charpentier

**Author notes:** Corresponding Author, Kathirvel Alagesan,; Max Planck Unit for the Science of Pathogens, Charitéplatz 1, 10117 Berlin, Germany.

## Abstract

Mass spectrometry is the premier tool for identifying and quantifying site-specific protein glycosylation globally. Analysis of intact glycopeptides often requires an enrichment step, after which the samples remain highly complex and exhibit a broad dynamic range of abundance.

Here, we evaluated the analytical benefits of high-field asymmetric waveform ion mobility spectrometry (FAIMS) coupled to nano-liquid chromatography mass spectrometry (nLC-MS) for analyses of intact glycopeptide devoid of any enrichment step. We compared the effects of compensation voltage on the transmission of N- and O-glycopeptides derived from heterogeneous protein mixtures using two FAIMS devices. We comprehensively demonstrate the performance characteristics of the FAIMS device for glycopeptide analysis and recommend optimal electrode temperature and compensation voltage (CV) settings for N- and O-glycopeptide analysis.

Under optimal CV settings, FAIMS-assisted gas-phase fractionation in conjunction with chromatographic reverse phase separation resulted in a 31% increase in the detection of both N- and O-glycopeptide compared to control experiments without FAIMS. Overall, our results demonstrate that FAIMS provides an alternative means to access glycopeptides without any enrichment providing an unbiased global glycoproteome landscape. In addition, our work provides the framework to verify ‘difficult-to-identify’ glycopeptide features.

## Introduction

Protein glycosylation is a highly dynamic co- and post-translational modification that can orchestrate a diverse range of biological functions, such as a biophysical and biochemical interface at the cell surface^1, 2^. System-wide MS analysis of linking glycans to peptides is a powerful analytical technique to investigate the structural-function relationship of glycoproteins. However, the robust and reliable identification of intact glycopeptides at the proteome scale remains technically challenging, owing mainly to the complexity and the macro- and microheterogeneity of site-specific glycosylation^3–5^. As such, glycopeptides are in low abundance compared to unmodified peptides after protease digestion. To tackle sample complexity and to alleviate issues associated with the co-elution of unmodified peptides, MS analysis of intact glycopeptide is dependent on various glycopeptide enrichment strategies such as HILIC, Lectin, etc^6^. Although highly valuable, these enrichment approaches have varying specificities for glycopeptides and may preferentially isolate sub-set of glycopeptides and add additional sample handling steps^7^. Another popular strategy is to leverage offline liquid chromatography (LC) fractionation (e.g., by strong-cation exchange or high-pH reversed-phase chromatography) to reduce the complexity, which can effectively increase the total number of identified glycopeptides. However, this step is time-consuming and laborious, requiring at least an additional day of sample preparation. For many PTMs, including glycosylation, acquiring enough material to perform these additional fractionation steps can also be challenging. Nonetheless, even after fractionation, the samples remain highly complex, exhibit a broad dynamic range of abundance, and require glycopeptide enrichment.

Field Asymmetric Waveform ion mobility spectrometry (FAIMS) separates gas-phase ions at atmospheric pressure based on their mobility through alternating high and low electric fields. Control of a direct current (DC) compensation voltage (CV) allows different groups of ions to pass through the FAIMS device to the MS. Previously, FAIMS has been successfully demonstrated to increase proteome coverage^8–10^, enrich modified peptides such as phosphopeptides^11^, simulated peptides, cross-linked peptides^12^ as well as bacterial glycopeptides^13^ and human N-glycopeptides^14^. Nevertheless, the use of FAIMS-nLC-MS/MS for simultaneous analysis of N- and O-glycopeptides has not been evaluated.

Here, we systematically explored the advantages and limitations of FAIMS as an alternate strategy for intact-glycopeptides independent of an enrichment step. First, we compared various CVs at fixed values to characterize the behavior pattern of glycopeptides utilizing two independent FAIMS Pro devices to access the universal applicability of the CVs specific for glycopeptides. Next, we compared the overall gain in glycopeptide identifications between conventional nLC-MS/MS and FAIMS-nLC-MS/MS with both fixed and sequential internal CV stepping. With these data, we recommend strategies for analyzing intact tryptic glycopeptides using FAIMS-nLC-MS/MS and comment on their suitability for N- and O-glycopeptide analysis. We also streamlined the glycoinformatics pipeline for the reliable and efficient analysis of glycopeptides. Ultimately, we demonstrate that using optimized gas-phase fractionation offered by FAIMS enables unbiased large-scale glycoproteomics, thereby enhancing both micro-and macro-heterogeneity.

## Materials and Methods

### Glycoprotein standard

The standard glycoprotein mixture consisted of four glycoproteins: Human Immunoglobulin G (IgG), Human Immunoglobulin M (IgM), Human Alpha-2-macroglobulin, and Bovine Fetuin. **Figure S1** represents known *N*- and *O*-glycosylation sites in these standard glycoproteins.

One hundred micrograms of individual glycoprotein standards were resuspended in 200 mM HEPES pH 8, 40 mM Tris(2-carboxyethyl)phosphine Hydrochloride (TCEP), 40 mM chloroacetamide (CAA) for reduction and alkylation of cysteine residues at 95 °C for 5 min. Prior to trypsin digestion, the samples were subjected to chloroform-methanol precipitation as described earlier^15^. The protein pellet was resolubilized in 50 mM TEAB buffer and trypsin was added in a 1:50 ratio (enzyme: substrate). After overnight incubation at 37 °C, the resulting glycopeptide/peptide mixtures were acidified with formic acid and dried in the speedvac without additional heating. After trypsin digestion, peptides were combined in equal parts by mass for the four proteins and used for the subsequent experiments.

### Reverse phase LC-MS/MS

All samples were resuspended in water containing 0.1 % Formic acid. Samples were separated by RP-HPLC using a Thermo Scientific Dionex UltiMate 3000 system coupled to either Thermo Scientific™ Orbitrap Exploris™ 480 or Orbitrap Fusion™ Lumos™ Tribrid™ Mass Spectrometer equipped with or without a FAIMS Pro device (Thermo Fisher Scientific) with Instrument Control Software version 3.3. Samples were loaded onto the PepMap C-18 trap-column (0.075 mm × 50 mm, 3 μm particle size, 100 Å pore size (Thermo Fischer Scientific)) 20 μL for 5 minutes with buffer A (0.1% formic acid, 2% acetonitrile) and separated over an in-house packed C18 column (column material: Poroshell 120 EC-C18, 2.7 μm (Agilent Technologies)) at 300 nL/min flow rate.

For FAIMS experiments, MS settings were identical except the FAIMS device was placed between the nano-electrospray source and the mass spectrometer. FAIMS separations were performed using the following settings: inner and outer electrode temperature = 100 °C (except where noted); Static FAIMS analyses were undertaken using CVs ranging from −10 to −120 V at 10 V increments. For standard glycoprotein mix, the nLC-MS/MS methods were 60 min total and each FAIMS condition was run in technical replicate. For reference serum proteins, the total run time was 60 min, and each LC-MS analysis was run in technical duplicates in Exploris and technical triplicates in Lumos. In the case of depleted human serum, the total run time was 120 min, and each condition was analyzed in technical duplicates. In all cases, a total of 350 ng peptide was injected per analysis.

All data were collected in positive ion mode using Data Dependent Acquisition mode. The ion source was held at 2200 V compared to the ground, and the temperature of the ion transfer tube was held at 275 °C. Survey scans of precursor ions (MS1) were collected in the mass-to-charge (*m/z*) range of 400 – 1800 at 60,000 resolution with a normalized AGC target of 200% and a maximum injection time of 25 ms, RF lens at 50%, data acquired in profile mode.

Monoisotopic precursor selection (MIPS) was enabled for peptide isotopic distributions, precursors of z = 2-7 were selected for data-dependent MS/MS scans for 3 seconds of cycle time for standard experiments (unless noted otherwise **see Table S1**) with an intensity threshold of 1.0e4. Dynamic exclusion was set to 40 seconds with a ±10 ppm window set around the precursor monoisotopic. Also, Precursor Fit was enabled for the MS/MS scans at the threshold of 70% within the 2 Da window.

MS2 scans were acquired using high-energy collision dissociation (HCD) fragmentation with a normalized collision energy of 30% with an isolation window of 2 Da, resolution of 15,000, AGC target of 200%, or a maximum injection time of 80 ms in a scan range determined automatically with the first mass set to 100 *m/z*. Detailed information on instrument-specific parameters is available in **Table S1**. To maximize instrument time, the presence of glycopeptide-specific oxonium ions (**Table S2**) in the scouting MS/MS scan was used to trigger an additional MS/MS scan ^16–18^ with a stepped collision energy HCD scan using NCE 15, 30, and 45% with a maximal injection time of 250 ms or AGC target of 500% at a resolution of 30,000 in an extended fixed scan range of 120-2000 Da.

### Data analysis

#### Byonic general parameters

All data were searched using Byonic software v3.10.10 (Protein Metrics) against the proteome database supplementary data with decoys appended within the Byonic environment as reversed sequences. Stepped FAIMS data files were processed within Byonic without splitting them into individual FAIMS CVs.

#### For standard glycoprotein mix

For all searches, cleavage specificity was set as semi-specific N-ragged for C-terminal to R and K residues and a maximum of two missed cleavage events was allowed. Carbamidomethyl (+57.021644) at cysteine was set as a fixed modification, oxidation of methionine (+15.994915) was included as a variable modification (common2), and acetylation at the protein N-terminus (+42.010565, rare1) as a variable modification. Gln -> pyro-Glu at N-terminal Q, Glu -> pyro-Glu at N-terminal E, and Ammonia-loss at N-terminal C were also allowed as variable modifications (rare1).

#### For human serum

In all cases, a total of three common and one rare modification were allowed per peptide/glycopeptide identification. A maximum precursor mass tolerance was set to 10 ppm with fragment mass tolerance set at 20 ppm. For glycopeptides, filtering metrics included a Byonic score greater than or equal to 200, and a logarithmic probability (logProb) value greater than or equal to 2. A glycan database containing 309 N-glycan compositions was used to identify intact N-glycopeptides. N-glycosylation was set as a variable modification (rare1). The O-glycan database used for O-glycopeptide searches consisted of 70 human O-glycans compositions. O-glycosylation was set as a variable modification (rare1).

## Results and Discussion

The FAIMS Pro interface has been reported to improve the signal-to-noise ratio, thereby increasing the detection of both modified and unmodified peptides when used as an orthogonal fractionation method^8,10, 19–23^. Although Zwitterionic Hydrophilic Interaction Liquid Chromatography (ZIC-HILIC) enrichment is considered unbiased, it has been demonstrated that the peptide sequences and the glycan size influence ZIC-HILIC glycopeptide enrichment efficiency^7^. Here, we sought to optimize and evaluate FAIMS as an alternate method for analyzing intact glycopeptides devoid of enrichment steps to improve unique glycopeptide identifications. Previously, FAIMS was successfully used to fractionate glycopeptides in gas-phase derived from bacteria^13, 24^ and human^14^. In this work, we expanded our analysis by testing samples of varying complexity at different LC gradient times using two independent FAIMS Pro devices coupled to either Exploris or Orbitrap Lumos MS (**Table S3**). We employed three samples of increasing complexity to generate a diverse set of glycopeptides. The first was a standard glycoprotein mix containing well-characterized glycoproteins **(Figure S1)**, the second was ERM-DA470k/IFCC reference human serum proteins containing the high background of albumin and finally, depleted human serum. Serum proteins were chosen as they represent an attractive source for clinical and diagnostic biomarkers^25^. Furthermore they display a high molecular complexity and dynamic range, and are also abundant in N- and O-glycoproteins^26–28^. In all cases, no specific glycopeptide enrichment step was performed.

### Optimal CV for N- and O-Glycopeptides

The inner and outer electrode temperature is known to influence FAIMS resolution^29^. First, we evaluated this phenomenon using tryptic glycopeptides derived from a standard protein mix analyzed at three independent inner electrode temperatures of 70, 85, or 100 °C, while the outer electrode was maintained at 100 °C. The resulting raw MS data were processed using Byonic for automated MS/MS-based (glyco)peptide identification. Reliable score-based glycopeptide identification largely depends on the MS fragmentation method^18^, the protein and glycan search space^30^, and glycopeptide characteristics such as peptide length, location of modification, or glycan composition^31^. Prior to evaluating the FAIMS performance for glycopeptide analysis, to ensure quality identifications, glycoPSMs returned from Byonic search were subjected to several filtering steps (***see methods for the sequence of post-Byonic search filtering steps-Figure S2***). We established two-step validation criteria for glycopeptides relying on glycan composition connectivity and glycoPSMs score.

First, we performed a simple yet effective visualization of glycan compositions using the GlyConnect Compozitor^32^ tool to correct the sparsity of missing compositions and identify potential outliers’ glycan clusters **(Figure S3)**. This step allowed us to confirm the *‘difficult-to-identify* glycopeptide features such as NeuAc, NeuGc, and multi-Fuc that are often mis-annotated^33^ and pass the empirical score. Next, we applied score and logProb cutoff to increase the confidence in glycopeptide identification. In this work, the glycan database used for glycopeptide identification is limited to the default glycan compositions available in Byonic. Nevertheless, visualization of glycan compositions via GlyConnect Compozitor^32^ enables the user to modify the glycan compositions and refine the search as required to reflect the context of the investigation. Notwithstanding, the logical next step would be to associate glycan composition-based visualization with every single glycopeptide detected to identify the outliers and increase confidence in glycopeptide identification. However, this goes beyond the scope of this investigation.

Applying the new filtering criteria, we observed an increase in the total number of glycopeptides with an increase in the inner electrode temperature. As such, diverse glycoforms are transmitted effectively for each peptide moiety, thereby increasing glycopeptide micro-heterogeneity, as shown in **Figure 1** and **Figure S4**. Our result also suggests that higher inner electrode temperature favors the transmission of highly charged species **(Figure S5)**. This feature is desirable for analyzing glycopeptides using EThCD or ETD fragmentation, enabling confident glyco-site localization^18, 31^.

**Figure 1:**
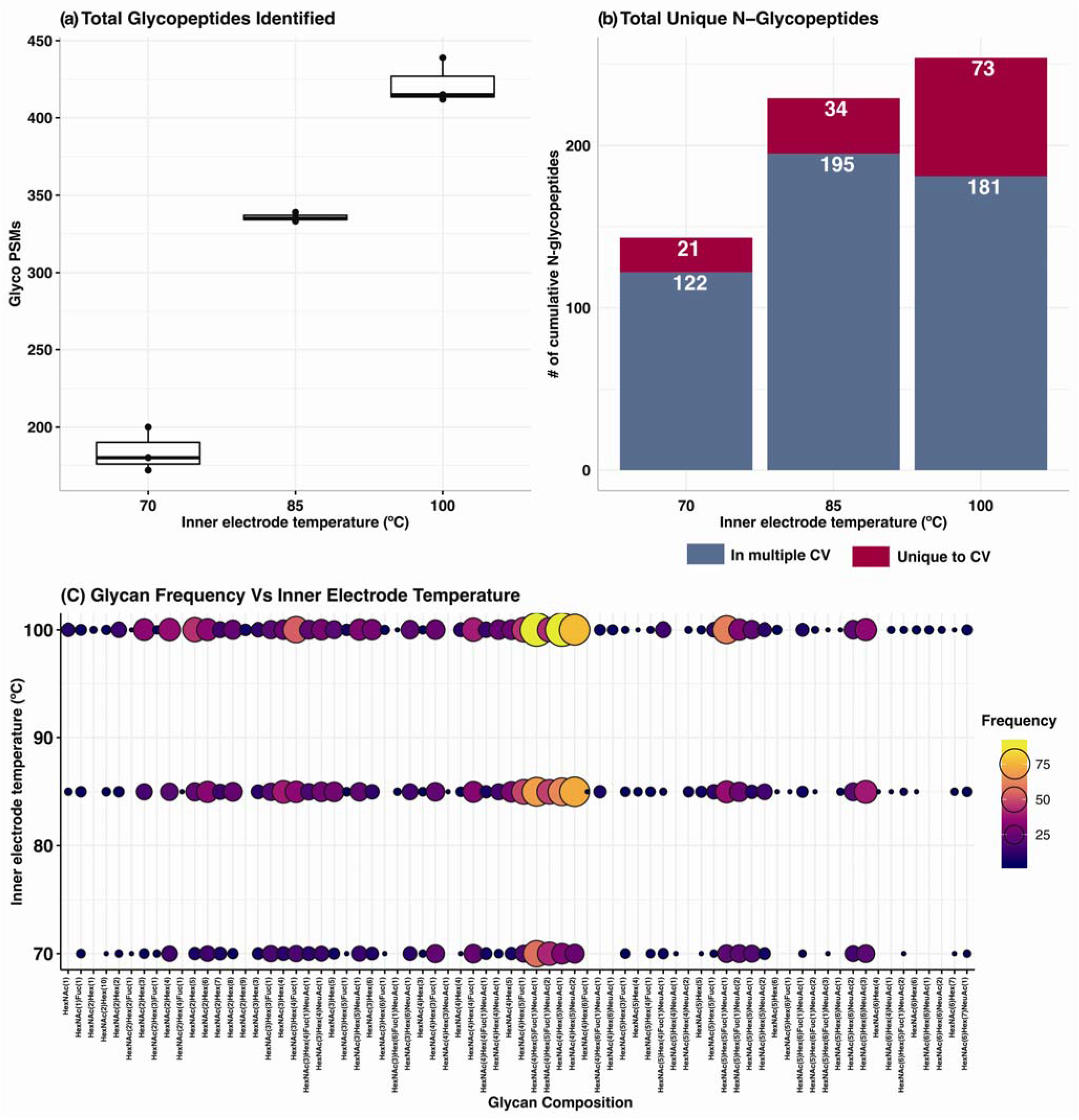
FAIMS inner electrode temperature effect on glycopeptide detection: (a) Total glycopeptides identified from triplicate analysis of tryptic peptides and glycopeptide mixture derived from standard protein at various FAIMS inner electrode temperature. (b) The stacked bar graph shows the cumulative number of unique N-glycopeptides identified with varying inner electrode temperatures. The blue bar indicates shared glycopeptides, whereas the burgundy bar indicates glycopeptides unique to the applied inner electrode temperature. An increase in inner electrode temperature improves the transmission of glycopeptides in FAIMS. In all cases, the outer electrode temperate was held at 100 °C. (c) Independent of peptide backbone length, an increase in inner electrode temperature allows for the efficient transmission of peptides carrying diverse glycan moieties. As such, higher inner and outer electrode temperature improves site-specific glycopeptide characterization, as shown in **Figure S4**

Having determined 100 °C as the optimal inner electrode temperature for glycopeptides, we then evaluated the transmission characteristics of glycopeptides in FAIMS by varying the applied CV. We performed LC-FAIMS-MS/MS experiments at individual CV ranging from −10 to −120 CV in 10 V increments. Each experiment was performed on a 350 ng injection of a standard glycoprotein tryptic digest using a 60 min gradient in triplicates. We compared the total glycopeptide identification (glycoPSMs) and the unique glycopeptides observed across this broad CV range. We detected a maximum number of unique glycopeptides within the CV range of −30 to −50, with a maximum at −40 (Exploris 480). No glycopeptides were observed above CV −90 **(Figure 2-a)**. Then, we fine-tuned the CV flanking the glycopeptide maxima with an increment of 5 V across the CV range of −25 to −80 V using tryptic peptides derived from the reference human serum. The unique N-glycopeptide identifications for each CV are shown in **Figure 2-b**. Irrespective of increased sample complexity, the CV distribution for the maximum number of unique glycopeptides remained comparable, with the highest unique N-glycopeptides identification rate at −45 V.

**Figure 2:**
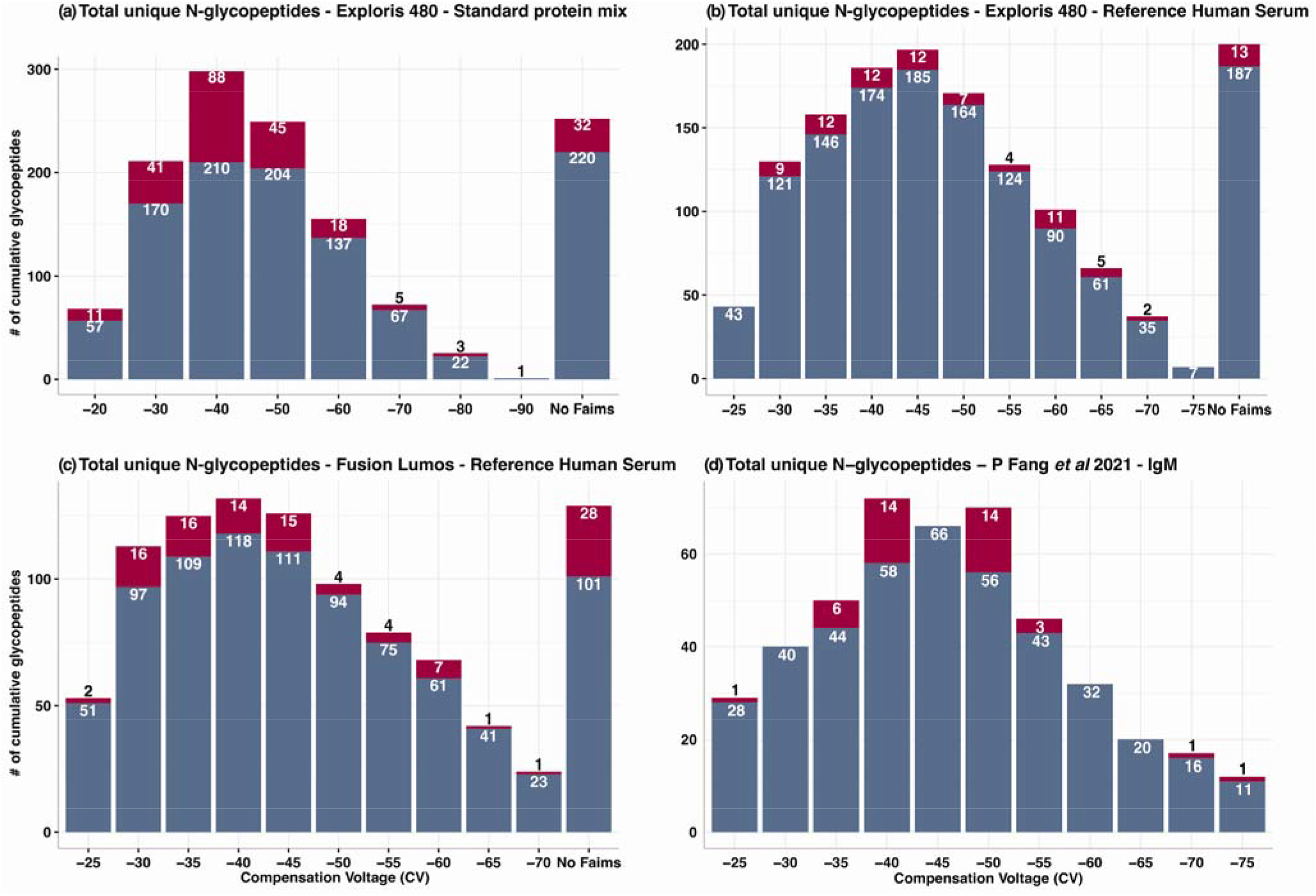
FAIMS enables gas-phase fraction of N-glycopeptides: (a-c) Distribution of N-glycopeptides across static CVs with varying sample complexity and two independent FAIMS devices coupled to either Exploris 480 or Lumos Fusion MS as indicated. All results are from 60 min analysis of trypsin digested protein source as indicated. Inner and outer electrode temperature was held at 100 °C. (a) standard protein mix, (b) ERM-DA470k/IFCC reference human serum proteins, (c) ERM-DA470k/IFCC reference human serum proteins, (d) reanalysis of Immunoglobulin M (IgM) glycopeptide data retrieved from PRIDE repository (PXD023951). The overlap and unique identification of glycopeptides across the CVs are represented in blue and burgundy, respectively. We observed that maximum transmission of N-glycopeptides with FAIMS occurs in the narrow CV range of −40 to −45.

In the reference human serum sample, the relative difference in the number of unique glycopeptides nLC-FAIMS-MS/MS & nLC–MS/MS analysis is underwhelming. 200 unique glycopeptides were identified without FAIMS compared to 197 at CV maxima (−45 V) **(*Figure 2-b & Table S3*)**. Since 65% of the total protein content is albumin in the reference human serum and FAIMS separation occurs after ionization, the stoichiometry of non-modified peptides is relatively higher than the standard protein mix. It is possible that despite optimal FAIMS CV voltage, lower ionization efficiency of glycopeptides might have hampered glycopeptide detection. As it is known that the presence of unmodified peptides suppresses glycopeptide ionization^34^. As previously noted, FAIMS is a continuous gas-phase filtration technique that allows for the detection of a subset of the ion population determined by the selected CV^19^. The cumulative number of unique N-glycopeptides identified within the CV range of −35 to −60 surpassed the analysis without FAIMS. In the case of Exploris 480, we identified a total of 232 unique N-glycopeptides across the CV range of −35 to −60 compared to 200 unique glycopeptides without FAIMS **(Figure 2b** and **Table S4)**, notwithstanding the increased instrumentation time required for the individual CV analysis. Overall, our results agree with a previous study^13^ and strongly indicate that FAIMS in combination with nLC-MS/MS is a useful *‘gas-phase enrichment’* technique providing access to glycopeptides that are otherwise not detected in the absence of FAIMS.

Recently, a broad CV range of −45 to −65 V was reported as an optimal CV for unlabeled glycopeptides with a maximum unique glycopeptide identification at two CVs: −45 V and −65 V^14^. Intrigued, we reanalyzed the IgM raw data from the PRIDE repository (PXD023951) using Byonic for automated glycopeptide identification **(See methods for search parameters)**. Contrary to the original report, we observed maximum unique glycopeptide identifications at −40 V, whereas −45 V resulted in the highest GlycoPSMs **(Figure 2-d)**. Additionally, we evaluated the glycopeptide behavior using a different FAIMS device coupled to OrbiTrap Lumos MS. As described earlier, we performed nLC-FAIMS-MS/MS experiments at individual CV ranging from −25 to −75 CV in 5 V increments in triplicates using tryptic peptides derived from the reference human serum. Here, consistent with our previous observation, unique N-glycopeptide identification maxima were observed at −40 V (**Figure 2-c**). Motivated by these results, we separately searched the reference serum protein dataset for O-glycopeptides, which is known to reduce false assignments^35^. In the case of O-glycopeptides, we observed CV maxima at −40 V and −35 V for Exploris and Fusion Lumos, respectively **(Figure S6)**. Taken together, our results suggest that each FAIMS Pro device is unique and requires CV optimization in the region of −30 to −50 V. We recommend 100 °C for both internal and external electrode temperature together with the CV of either −40 or −45 V as a starting point to identify the *‘sweet spot’* unique to the FAIMS device to maximize the glycopeptide identification **(Figure 2 a-d)**.

### Internal stepped CV for N- and O-Glycopeptides

Using multiple CVs within a single analysis run allows for continuous gas-phase fractions resulting in higher identification of peptides. Having established the optimal CV for glycopeptides, we evaluated whether increasing the number of CVs within a single analysis run (Internal stepped CV) increases the unique glycopeptide identifications. To this end, we combined the CVs in 10 and 15 V steps centered around the CV that provided maximum identification (CV = −40 V for O-glycopeptides or −45 V for N-glycopeptides – **Figure S6 & Figure 2**).

The cumulative number of unique glycopeptide identifications from two individual runs obtained for each CV combination indicates that FAIMS-nLC-MS/MS-based glycopeptide analysis outperformed standard nLC-MS/MS-based analysis (**Figure 3 a-b**). Of all two CV combinations tested, the −40/−50 V combination resulted in the higher identification rate for both N- and O-glycopeptides in Exploris. Under No-FAIMS conditions, we identified 356 unique N-glycopeptides, whereas the −40/−50 V combination resulted in 469 unique N-glycopeptides. In the case of O-glycopeptides, we identified 326 unique O-glycopeptides with FAIMS at CV combination −40/−50 V in contrast to 246 without FAIMS. Note that the overall total GlycoPSM identified in nLC-MS/MS condition is relatively higher than all two CV combinations tested. In parallel, we also performed the single CV analysis for the CV range −35 to −55 V **(*Figure 3 c-d*)**. We observed that the CV optima for glycopeptides remain stable irrespective of sample complexity and gradient length for N-glycopeptides at −45 V. In the case of O-glycopeptides, we observed that sample complexity influences the CV maxima to a small extent. Previously, when analyzing reference serum, the CV maxima for O-glycopeptide was at −40 V and for the depleted human serum it was −45 V. Comparing analysis without FAIMS to FAIMS with two CV combinations ±5 V around CV maxima resulted in higher identifications of both N- and O-glycopeptide **(Figure 3 a-b and S7)**.

**Figure 3:**
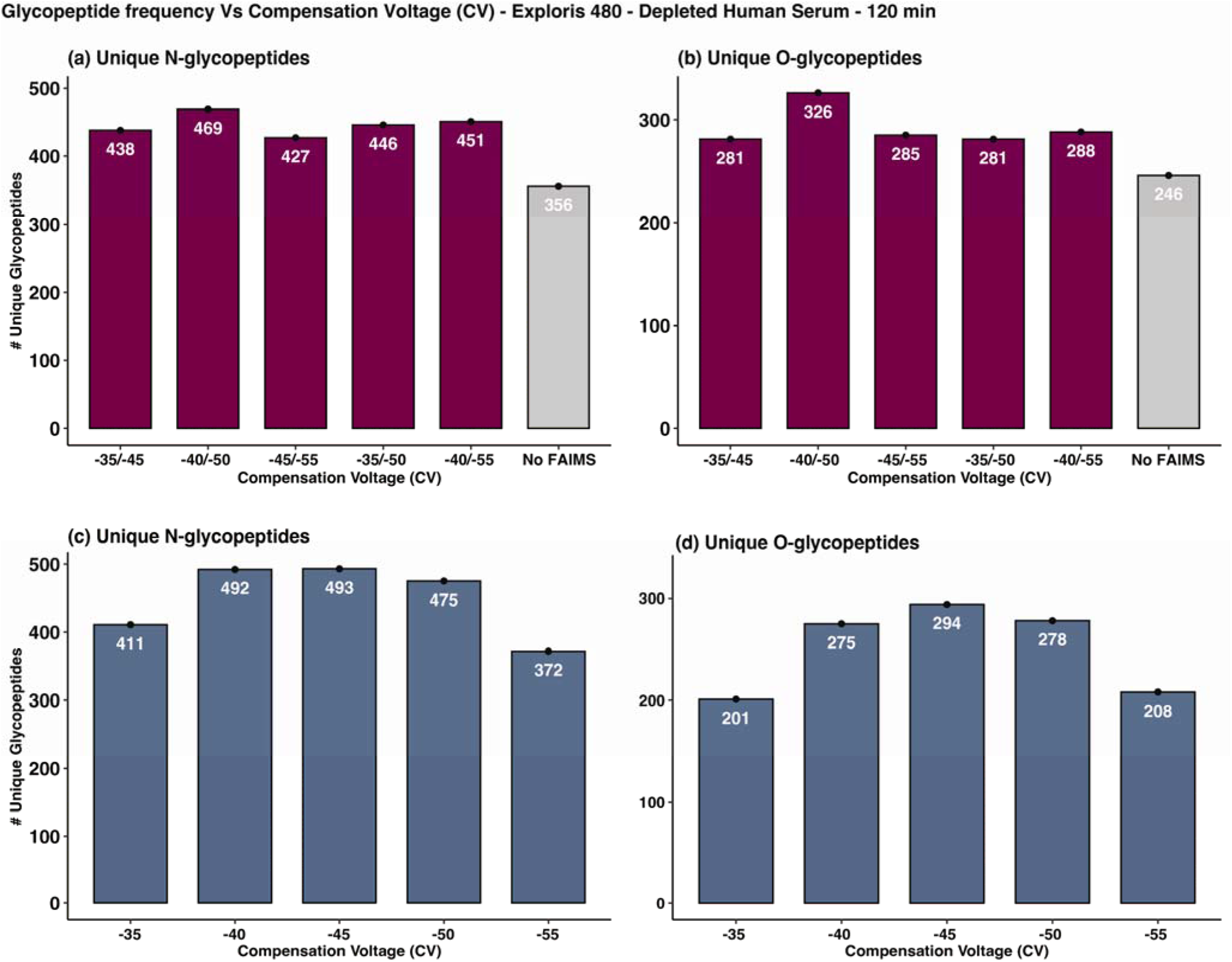
Optimization of combining multiple CVs for bottom-up glycoproteomics: All data are from the duplicate analysis of trypsin digested depleted human serum within 120 min analytical run (a) Number of unique N-glycopeptides (b) Number of unique O-glycopeptides identified under identical nLC-MS/MS with No FAIMS and FAIMS two CV combination as indicated. As shown, stepped CV FAIMS outperformed No FAIMS analysis demonstrating the superior potential of FAIMS to enrich for both N- and O-glycopeptides in the gas phase under identical nLC-MS/MS conditions. Number of unique glycopeptides identified under static CV voltage −35 V to −55 V (c) for N-glycopeptides (d) for O-glycopeptides.

Our results demonstrate that FAIMS can effectively ‘enrich’ both N- and O-glycopeptides in the gas-phase under the same CV conditions. When combining two CVs in 10 V intervals around the apex CV, excellent depth for both N- and O-glycopeptide was achieved (Figure 3 and S7). Further improvements in acquisition strategies, such as a global dynamic exclusion parameter for complete analysis rather than for each CV experiment, especially during CV switching, can increase the number of unique glycopeptides by minimizing the overlap during the CV switch.

## Conclusion

Selective enrichment of glycopeptides is essential for intact MS analysis of glycopeptides to compensate for their low stoichiometry due to microheterogeneity. As such, glycopeptides exhibit reduced ionization efficiency compared to non-modified peptides in MS detection. In this study, we evaluated the analytical benefits of FAIMS coupled with nLC-MS/MS as an alternative strategy to improve glycopeptides detection devoid of any enrichment step. Utilizing a sample of varying complexity, we demonstrate that FAIMS-assisted glycopeptide analysis enhances the detection of both N- and O-glycopeptides. In a head-to-head comparison of tryptic glycopeptides derived from depleted serum by nLC-MS/MS with and without FAIMS, we observed that the optimal CV setting resulted in a 31% increase in both N- and O-glycopeptides identified **(Figure 3 a-b)**. Our data suggest that FAIMS-assisted analysis significantly maximizes the depth of glycopeptide coverage by improving the site-specific glycosylation profile. We also introduced a glycan composition filtering step prior to the score cutoff to increase confidence in the glycopeptide assignment.

It should be noted that optimal CV is dependent on the individual FAIMS device. Nonetheless, a similar trend exists in glycopeptide transmission with applied CV with apex identification at CV −40 and −45 V for two independent FAIMS devices. In summary, we believe that the FAIMS-assisted bottom-up glycoproteomics workflow improves glycopeptide analysis without any enrichment step.

## Supporting information

proteome database

proteome database

## Acknowledgments

This work was supported by the Max Planck Society. We thank Florian Kondrot for his excellent technical support.

## Conflict of interest

None

## Author Contributions

***K.A*** – Conceptualization, Methodology, Software, Validation, Investigation, Formal analysis, Data curation, Writing – Original draft, Writing – Review & Editing, Visualization, Project administration.

***R.A.B*** - Data curation, Visualization, Writing – Review & Editing.

***E.C*** - Resources, Writing – Review & Editing, Funding acquisition.

The manuscript was written with the contributions of all authors. All authors have approved the final version of the manuscript.

## Data availability

The mass spectrometry proteomics data have been deposited to the ProteomeXchange Consortium via the PRIDE partner repository. Data are available via ProteomeXchange with identifier PXD038673. List of identified glycopeptides are provided as Supplementary Data.

Reviewer account details:

Username

Password: 30KsTXRC

## Notes

### Competing Interest Statement

The authors have declared no competing interest.

