## Supplementary Data for "Improved N- and O-Glycopeptide Identification using High-Field Asymmetric Waveform Ion Mobility Spectrometry (FAIMS)"

### Author information

#### ORCiD

##### KA - 0000-0002-7596-5558

RAB	-	0000-0002-0656-1795

##### EC - 0000-0002-0254-0778

#### *Corresponding Author

Kathirvel Alagesan,; Max Planck Unit for the Science of Pathogens, Charitéplatz 1, 10117 Berlin, Germany.

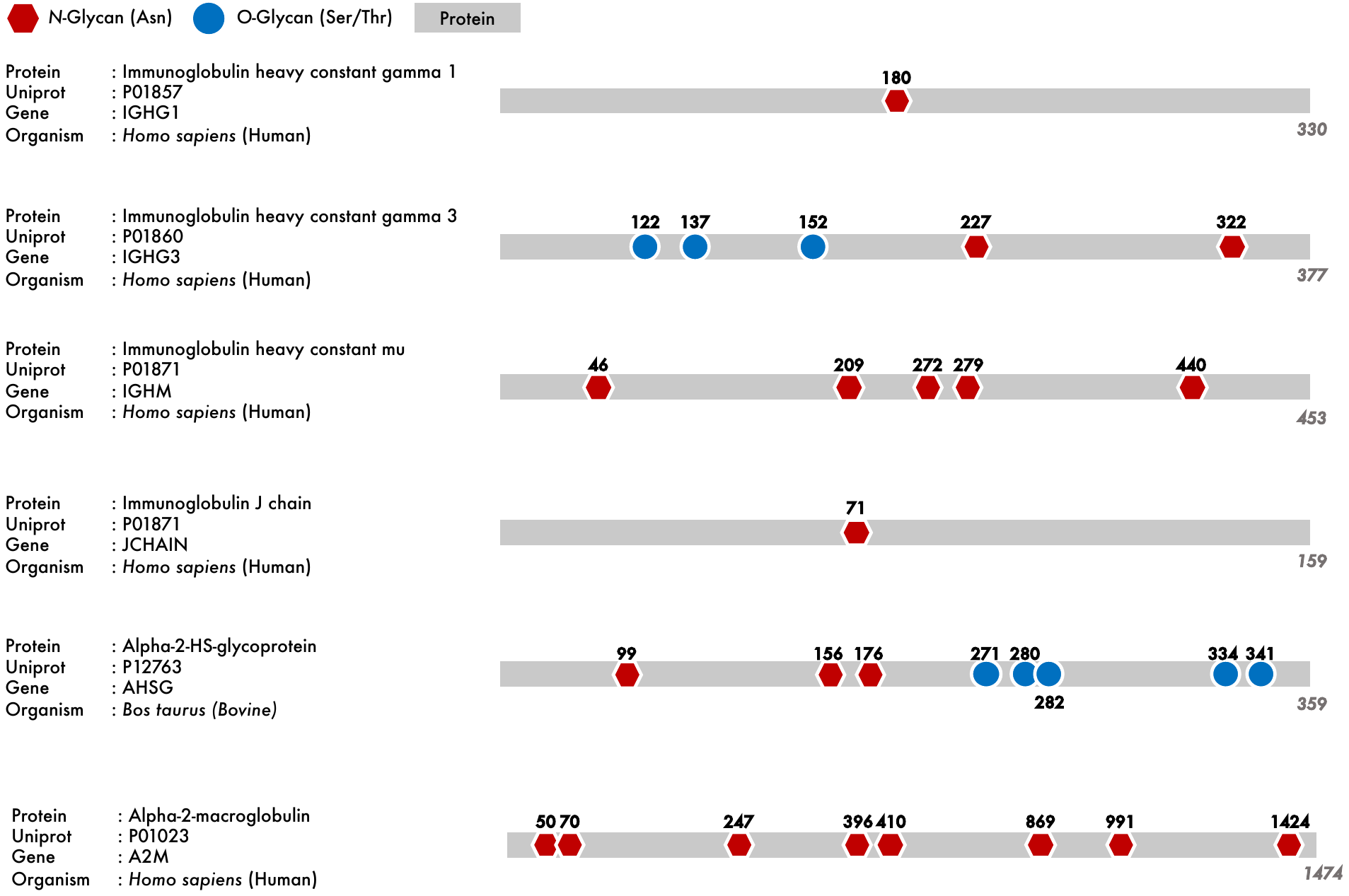

Figure S1: Standard glycoproteins used in this study. The number of known *N*- and *O*-glycosylation sites available for each standard protein in Uniprot are shown in red hexagon and blue circle, respectively. These proteins were chosen as they are well characterized and known to generate both *N-* and *O-*glycopeptides with varying degrees of glycan complexity and peptide lengths. Furthermore, glycopeptides derived from these glycoproteins after trypsin digestion were subjected to sequential exoglycosidase digestion to further diversify the glycopeptide repertoire.

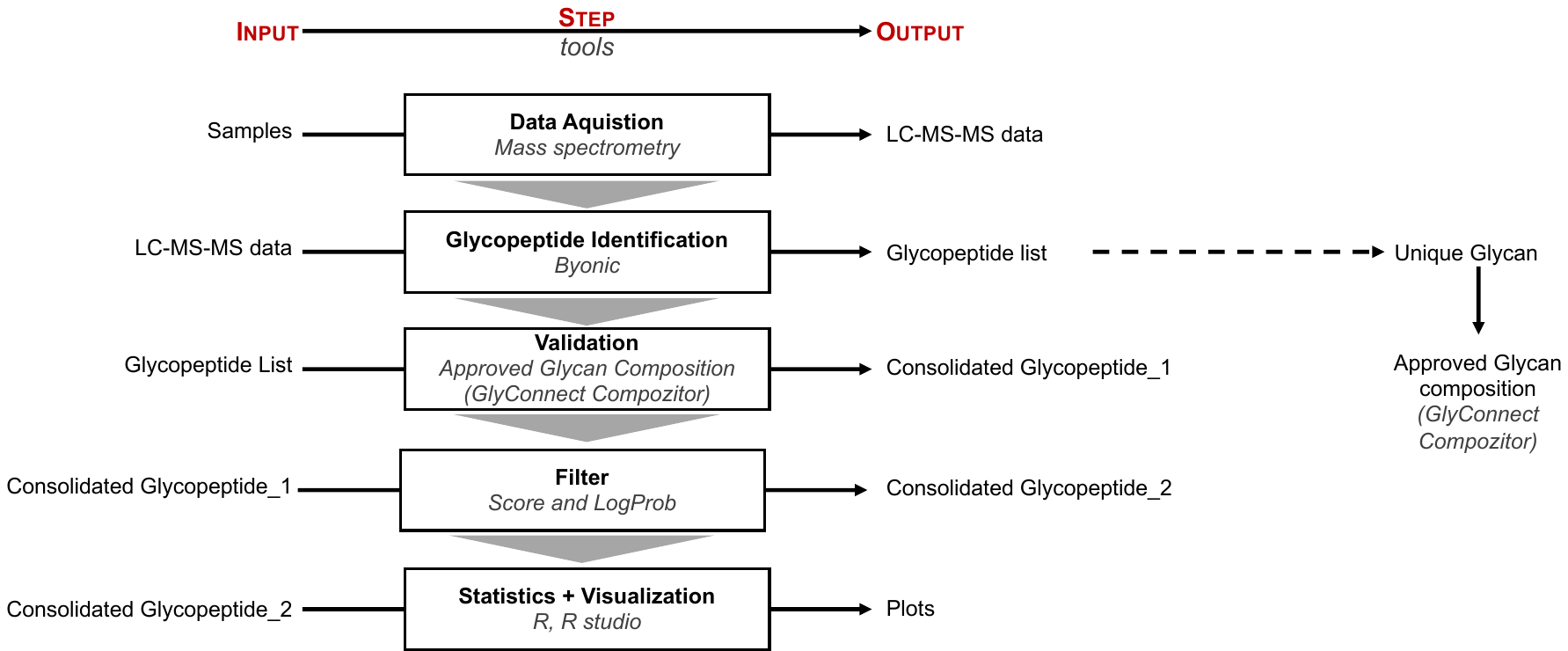

**Figure S2:** Sequence of glycopeptide data analysis and visualization steps. Byonic was used for automated MS/MS-based (glyco)peptide identification. In the traditional analysis, the Byonic score and log probability (Log Prob) score are used as criteria to ensure confident glycopeptide identification. However, this step does not exclude any potential misassignments of glycan composition, especially when dealing with difficult-to-identify glycopeptide features. To address this, we included a “Glycan composition” based visualization feature using the Glyconnect compozitor tool. This additional step allows for the visualization of glycan pathways based on the compositions. As we utilize the cumulative glycopeptide list from all experimental conditions to extract glycan composition, we minimize the number of virtual nodes (missing glycan compositions) observed in the filtering step. This step enables the identification of outliers easily. In the next step, the cumulative glycopeptide is filtered based on “approved glycan composition” as exported from the Glyconnect compozitor tool.

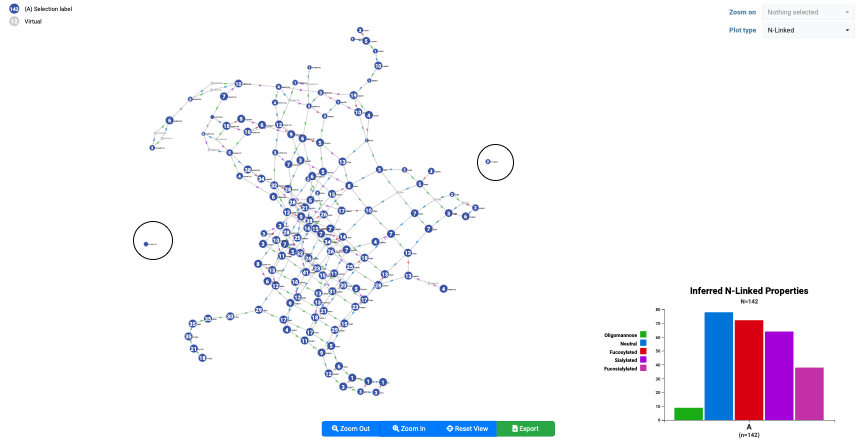

**Figure S3:** Exemplary glycan composition visualization using the Glyconnect compozitor tool. Unique glycan composition retrieved from the cumulative list of glycopeptides is visualized here. On a quick look, we can identify two outlier glycan compositions circled in black as they do not have any connections to other glycan compositions. Upon further evaluation, we can export the selected list of glycans from the tool to further filter the glycopeptides based on glycan composition and proceed further with the score and log prop based filtering.

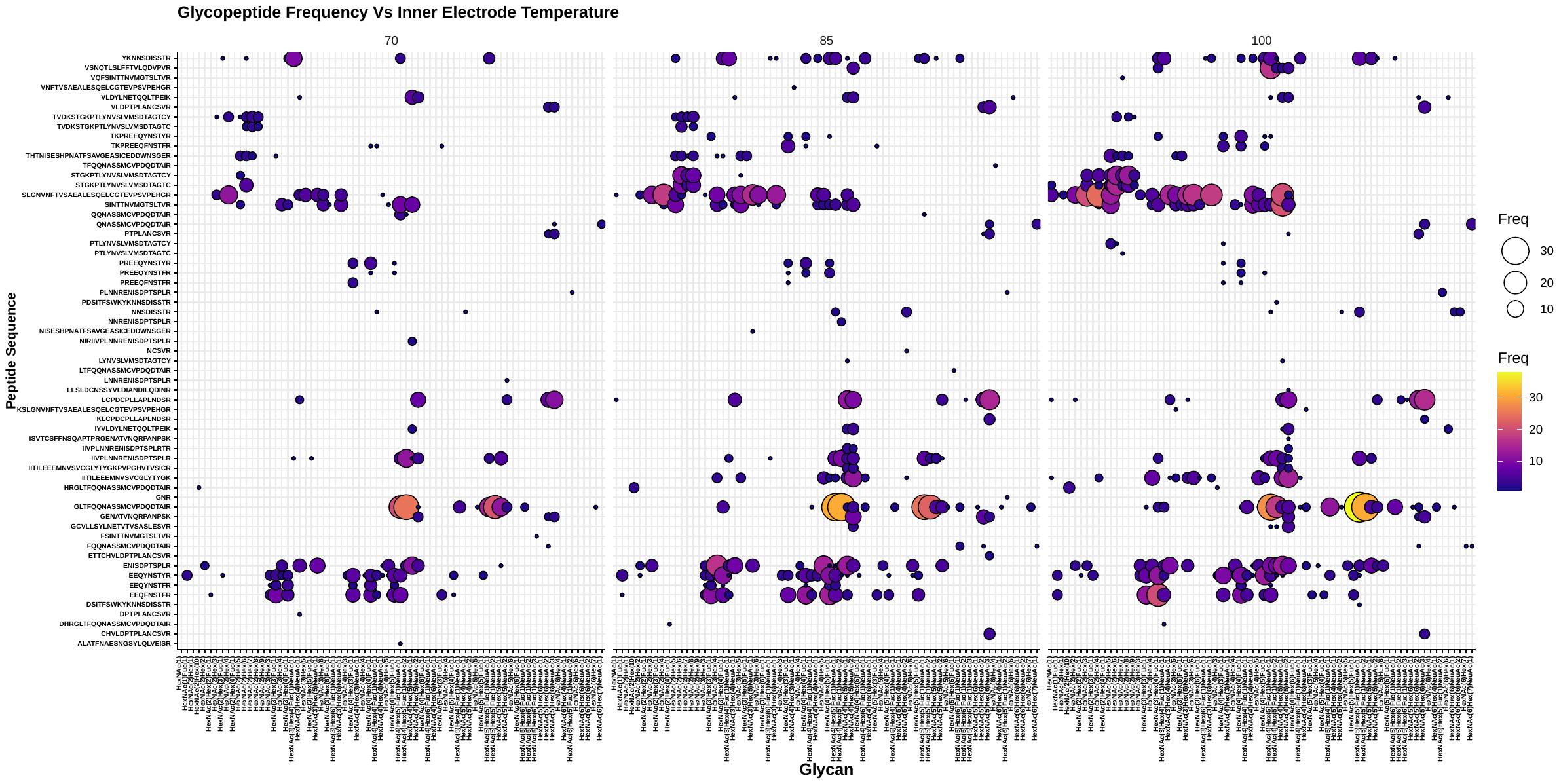

**Figure S4: Influence of inner electrode temperature on glycopeptide microheterogeneity. Increase in inner electrode temperature improves the transmission of glycopeptide carrying diverse glycan moieties thereby improving site-specific glycopeptide characterization.**

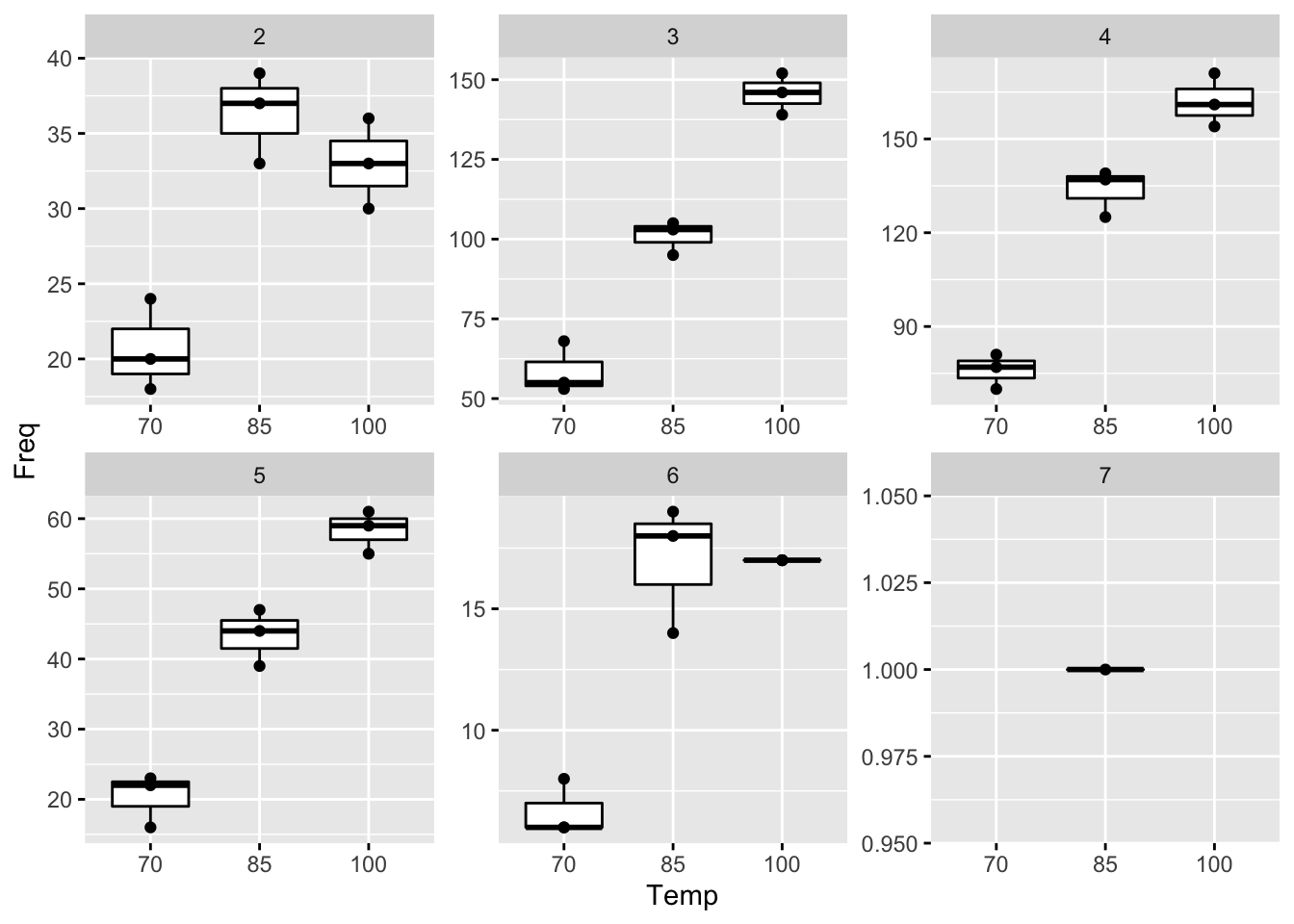

**Figure S5:** Effect of reducing the inner electrode temperature on the precursor charge state of glycopeptides identified. The outer electrode was held constant at 100 °C. These charge-state specific distributions of glycopeptides indicate that FAIMS can be used to preselect populations of glycopeptides with charge states optimal for a given fragmentation method.

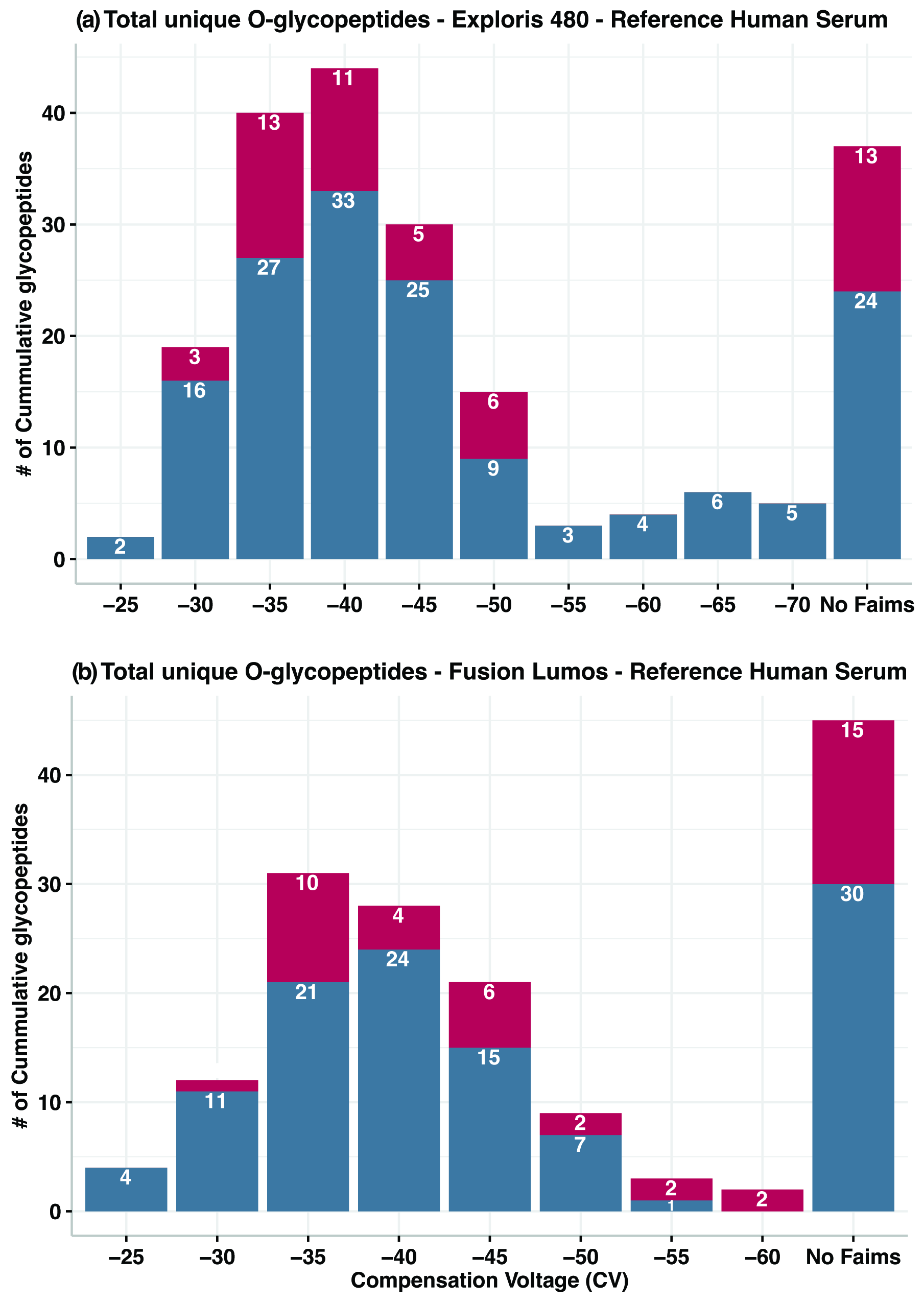

**Figure S6:** **FAIMS enables gas phase fraction of O-glycopeptides.** Distribution of O-glycopeptides across static CVs using two independent FAIMS devices coupled to either Exploris 480 or Lumos Fusion MS as indicated. All results are from 60 min analysis of trypsin-digested reference human serum proteins.

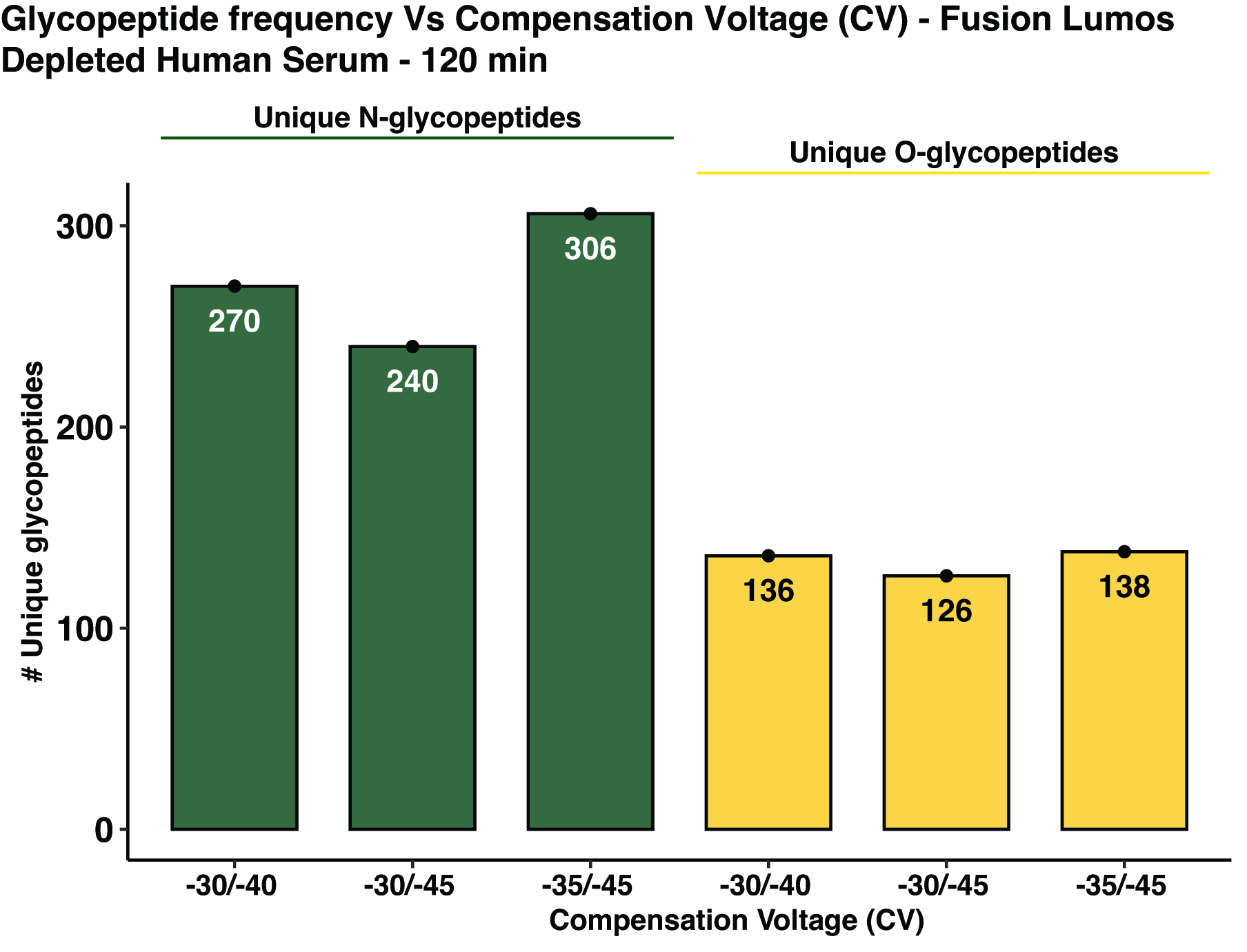

**Figure S7: Optimization of combining multiple CVs for bottom-up glycoproteomics in Fusion Lumos.** Number of unique *N-* and *O-*glycopeptides identified in two CV combination. All data are from duplicate analysis of trypsin-digested depleted human serum within a 120 min analytical run.

**Table S1:** MS and MS/MS acquisition parameters for the analysis of intact glycopeptides using Exploris 480 and Orbitrap Fusion Lumos Tribrid Mass Spectrometer.

| **Parameter** | **Exploris 480** | **Orbitrap Fusion Lumos** |
| --- | --- | --- |
| Ionization mode | Positive | Positive |
| Electrospray voltage | 2200 V | 2200 V |
| MS1 resolution | 60 K | 60 K |
| MS1 scan range | 400 – 1800 | 350 – 1800 |
| MS1 maximum injection time (ms) | 50 | 50 |
| MS1 normalized AGC target | 200 | 250 |
| MS2 resolution | 15 k | 15 k |
| MS2 scan range | Auto with the first mass set to 100 m/z | Auto with the first mass set to 100 m/z |
| MS2 maximum injection time (ms) | 60 | 60 |
| MS2 normalized AGC target | 120 | 120 |
| Isolation | 2 *m/z* with quadrupole | 2 *m/z* with quadrupole |
| Fragmentation (Normalized) | HCD at 30% | HCD at 35% |
| Dynamic exclusion | 40 with a ±10 ppm window | 40 with a ±10 ppm window |
| Charge states | 2–7 selected | 2–7 selected |
| Intensity threshold | 5.00E+03 | 2.5E+04 |
| DDA method | 3 s cycle time* | 3 s cycle time* |
|  | **Product dependent scan#** | |
| Fragmentation | sHCD | sHCD |
|  | NCE 20, 35, and 50% | NCE 20, 35, and 50% |
| MS2 resolution | 30 K | 30 K |
| MS2 scan range | 120-2000 Da | 110-2000 Da |
| MS2 normalized AGC target | 500% | 500% |
| MS2 maximum injection time | 250 | 250 |
|  | # See Table S2 for the list of product ions for product dependent scans | |
|  | *see below for cycle time for internal CV stepping | |

**CV and cycle time design for internal CV stepping for MS analysis of depleted human serum**

| **Experiment Type** | **Number of CVs** | **Number of Experiments** | **CVs (V)** | **Time/MS experiment (s)** | **Total Cycle Time (s)** |
| --- | --- | --- | --- | --- | --- |
| Control | NA | 1 | NA | NA | 4 |
| Control | NA | 2 | NA | 2 | 4 |
| FAIMS | 2 | 2 | -35/-45 | 2 | 4 |
| FAIMS | 2 | 2 | -40/-50 | 2 | 4 |
| FAIMS | 2 | 2 | -45/-55 | 2 | 4 |
| FAIMS | 2 | 2 | -35/-50 | 2 | 4 |
| FAIMS | 2 | 2 | -40/-55 | 2 | 4 |
| FAIMS | 2 | 2 | -30/-40 | 2 | 4 |
| FAIMS | 2 | 2 | -30/-45 | 2 | 4 |

**Table S2:** List of glycopeptide oxonium ions used for product dependent MS/MS scan to maximize glycopeptide identifications.

| **Sugar Composition** | **Diagnostic Oxonium Ion**  **(*m/z*)** | **Diagnostic Oxonium Ion – H_2_O**  **(m/z)** |
| --- | --- | --- |
| HexNAc | 204.0867 | 186.0761 |
| Hex | 163.0601 | 145.0495 |
| Hex-HexNAc | 366.1395 | - |
| Hex-HexNAc-Hex | 528.1923 | - |
| NeuAc | 292.1027 | 274.0921 |
| NeuAc-Hex-HexNAc | 657.2349 | - |
| NeuAc-Hex-HexNAc-Hex | 819.2877 | - |
| dHex | 147.0652 | 129.0546 |
| Hex-HexNAc-dHex | 512.1974 | - |

**Table S3:** Overview of samples analyzed.

| **Sample Type** | **Gradient Length (min)** | **Instrument** | **Condition** |
| --- | --- | --- | --- |
| Standard glycoprotein mix | 60 | Exploris 480 | Variable Inner Electrode Temperature: 70, 85 and 100 °C |
| Standard glycoprotein mix | 60 | Exploris 480 | Varying FAIMS CV Voltage: -10 to -120 CV in 10 V |
| Reference Human serum | 60 | Exploris 480 | Varying FAIMS CV Voltage: -25 to -80 V CV in 5 V increments |
| Reference Human serum | 60 | Fusion Lumos | Varying FAIMS CV Voltage: -25 to -80 V CV in 5 V increments |
| Reference Human serum | 120 | Exploris 480 | Varying FAIMS: -35, -40, -45, -50, -55  Internal Stepping CV: -35/-45, -40/-50, -45/-55, -35/-50, -40/-55 |
| Reference Human serum | 120 | Fusion Lumos | Internal Stepping CV: -30/-40, -30/-45, -35/-45 |

**Table S4:** Overview of unique N-glycopeptides identified at CV maxima and region flanking CV maxima.

| **MS** | **Sample** | **nLC-MS/MS** | **nLC-FAIMS-MS/MS** | |
| --- | --- | --- | --- | --- |
| **Exploris 480** | Standard Glycoprotein Mix | 252 | 298 | CV= -40 |
|  | Reference Serum | 200 | 197 | CV= -45 |
|  | Reference Serum | 200 | 232 | CV = -35 to -60 |
| **Fusion Lumos** | Reference Serum | 129 | 132 | CV=-40 |
|  | Reference Serum | 129 | 183 | CV = -30 to -50 |

#### Description of Additional Supplementary Files

**Supplementary Data 1:** Fig_1_Souredata_InnerElectrode_Glycopeptides.csv
Description: Information about identified glycopeptides from varying inner electrode temperature.

**Supplementary Data 2:** Fig2a_N_Glycopeptides_StdProteinMix.csv
Description: Information about identified N-glycopeptides derived from standard protein mix from varying CV (Exploris 480).

**Supplementary Data 3:** Fig_2b_N_Glycopeptides_RefHumanSerum.csv
Description: Information about identified N-glycopeptides derived from reference human serum (Exploris 480) from varying FAIMS CV.

**Supplementary Data 4:** Fig_2c_N_Glycopeptides_RefHumanSerum.csv
Description: Information about identified N-glycopeptides derived from reference human serum (Fusion Lumos) from varying FAIMS CV.

**Supplementary Data 5:** Fig_2d_IgM_N_Glycopeptide.csv
Description: Information about identified glycopeptides derived from IgM P Fang et al 2021.

**Supplementary Data 6:** Fig_3a_c_N_Glycopeptides_DepletedHumanSerum.csv
Description: Information about identified N-glycopeptides from static and internal stepped CV from depleted human serum (Exploris 480).

**Supplementary Data 7:** Fig_3b_d_O_Glycopeptides_DepletedHumanserum
Description: Information about identified O-glycopeptides from static and internal stepped CV from depleted human serum (Exploris 480).

**Supplementary Data 8:** Fig_S6_O_Glycopeptides_RefHumanSerum.csv

Description: Information about identified O-glycopeptides from static CV from depleted human serum (Exploris 480).

**Supplementary Data 9:** O_Glycopeptides_Lumos_DepletedHumanSerum_sFig7.csv

Description: Information about identified O-glycopeptides from internal stepped CV from depleted human serum (Fusion Lumos).

**Supplementary Data 10:** GlycoFocusedProteinDB.txt
Description: Fasta file that has proteins identified in the proteomics experiment. This database was used to search intact glycopeptide spectra for the reference human serum.

**Supplementary Data 10:** uniprot-proteome_UP000005640_Human.txt

Description: Fasta database used to search intact N- and O-glycopeptide spectra for the depleted human serum
